# ORFeus: A Computational Method to Detect Programmed Ribosomal Frameshifts and Other Non-Canonical Translation Events

**DOI:** 10.1101/2023.04.24.538127

**Authors:** Mary O. Richardson, Sean R. Eddy

## Abstract

**Background:** Canonical protein translation requires that ribosomes initiate translation at the correct start codon, maintain a single reading frame throughout elongation, and terminate at the first in-frame stop codon. However, ribosomal behavior can deviate at each of these steps, sometimes in a programmed manner. Certain mRNAs contain sequence and structural elements that cause ribosomes to begin translation at non-canonical start codons, shift reading frame, read through stop codons, or reinitiate on the same mRNA. These processes represent important translational control mechanisms that can allow an mRNA to encode multiple functional protein products or regulate protein expression. The prevalence of these events remains uncertain, due to the difficulty of systematic detection.

**Results:** We have developed a computational model to infer non-canonical translation events from ribosome profiling data.

**Conclusion:** ORFeus identifies known examples of alternative open reading frames and recoding events across different organisms and enables transcriptome-wide searches for novel events.

## Background

Deviation from the rules of canonical protein translation can lead to protein synthesis from alternative ORFs (altORFs). Altered initiation may result in upstream ORFs (uORFs) or downstream ORFs (dORFs) [1, 2]. Elongation can be affected by programmed ribosomal frameshifting (PRF), where the ribosome slips forward or backward (usually by +1 or -1 nucleotide) and changes reading frame during translation [3, 4]. Termination may be overridden by stop codon readthrough (SCR) or incorporation of selenocysteine or pyrrolysine, which lead to extended translation past an in-frame stop codon [5, 6]. These non-canonical translation events generate alternate protein sequences and are an important feature of the translational landscape of many organisms. Alternative translation events, especially frameshifts, are common in viruses [7], but alternate translation mechanisms are also observed beyond viruses, from bacteria to humans. It is increasingly clear that these events may be more widespread than we realize, with known examples spanning all domains of life [8–17]. Non-canonical events can act as a regulatory switch for protein synthesis or can enable synthesis of multiple different functional protein products from a single mRNA. Together, frameshifts and other non-canonical events play an important role in regulating translation and producing alternate peptides that may have gone undetected using standard annotation pipelines.

One method for detecting these alternate translation events is ribosome profiling (ribo-seq). Ribo-seq is a technique that provides nucleotide-resolution information about ribosome position during translation, which can be used to infer open reading frames (ORFs). The protocol for ribo-seq involves (i) treating ribosome-bound mRNAs with a nuclease – typically RNase I or micrococcal nuclease (MNase), (ii) isolating the ribosome-protected fragments (or ‘footprints’) (iii) generating a library for deep sequencing, and (iv) mapping the ribosome footprints back to the genome or transcriptome [18]. The pattern of mapped reads can then be used to identify translated regions by manually or computationally searching for regions of high read density. In many ribo-seq data sets (though not all), aggregate ribosome footprint density over coding genes shows a characteristic triplet periodicity. This periodicity suggests that ribo-seq data can not only be used to measure ribosome density in an ORF at course resolution, but also to measure the reading frame that ribosomes are in at nucleotide resolution.

However, this requires looking at the data one ORF at a time, and at the single-ORF level the signal is often much more sparse. While some nucleotide (nt) positions have a high footprint density, the majority of positions in an mRNA typically have no mapped footprints. This heterogeneity presents a challenge for directly inferring frame at each nt position of a gene. Ribo-seq data can also exhibit a high level of noise, due in part to nuclease sequence bias and variability in footprint length [19]. This is especially pronounced in initial work in bacteria, where the wide range of footprint lengths blurs the signal and makes it difficult to decipher which nucleotide is in the P-site. However, subsequent work has demonstrated that adding the endonuclease RelE (in addition to MNase) during ribosome profiling in *E. coli* results in clear triplet periodicity of footprint ends (in aggregated metagenes) [20]. Additionally, computational approaches have been developed to determine the position of the P-site within different length ribo-seq fragments, which improves resolution for data sets generated with MNase, RNase I, or other nucleases [21].

Numerous methods exist to detect ORFs from ribo-seq data based on the distribution, periodicity, or read lengths of footprints in actively translated regions. Many of these algorithms allow for detection of novel ORFs, alternative initiation, and short uORFs or dORFs [22–33]. Some allow for ORFs with non-AUG start codons. However, current methods fall short when searching for recoding events that break the rules of canonical translation. Recoding events like programmed ribosomal frameshifts and stop codon readthrough violate the assumptions of current models, making detection difficult. Ribo-seq data has the power to reveal these recoding events [16, 34–36], but there is no integrated approach available. Incorporating detection of all types of alternative ORFs (altORFs) and recoding events into a single method would allow for a more complete annotation process and help flag unexpected translation events for investigation.

Here we present ORFeus, a general-purpose computational tool for inferring altORFs and recoding events. ORFeus uses a hidden Markov model to infer translation patterns from ribo-seq data that is inherently noisy and sparse. The model identifies changes in reading frame and additional upstream or downstream reading frames, making it suitable for detection of many alternative translation events. Given high coverage, periodic ribo-seq data, ORFeus can identify novel or extended ORFs (including uORFs and dORFs) with either canonical or non-canonical start codons, as well as programmed ribosomal frameshifts and stop codon readthrough events.

## Results

### Data processing

ORFeus takes as input aligned ribosome profiling data, reference annotations, and a reference genome sequence. The annotations file should contain known 5’UTR, 3’UTR, and protein-coding ORF features (although for bacteria, UTRs are typically not annotated). Aligned ribo-seq reads submitted to ORFeus should be uniquely mapped to the genome and have their 5’ ends (or 3’ ends) offset to correspond to the P-site of the ribosome (see Methods for further explanation) (Fig. 1A). Mapping reads uniquely to the genome is advised (though not strictly required) to avoid confounding signal from multimapped reads that may be mistaken for alternative translation. Available tools for alignment and pre-processing include RiboGalaxy [37] and Shoelaces [21].

**Fig. 1.**
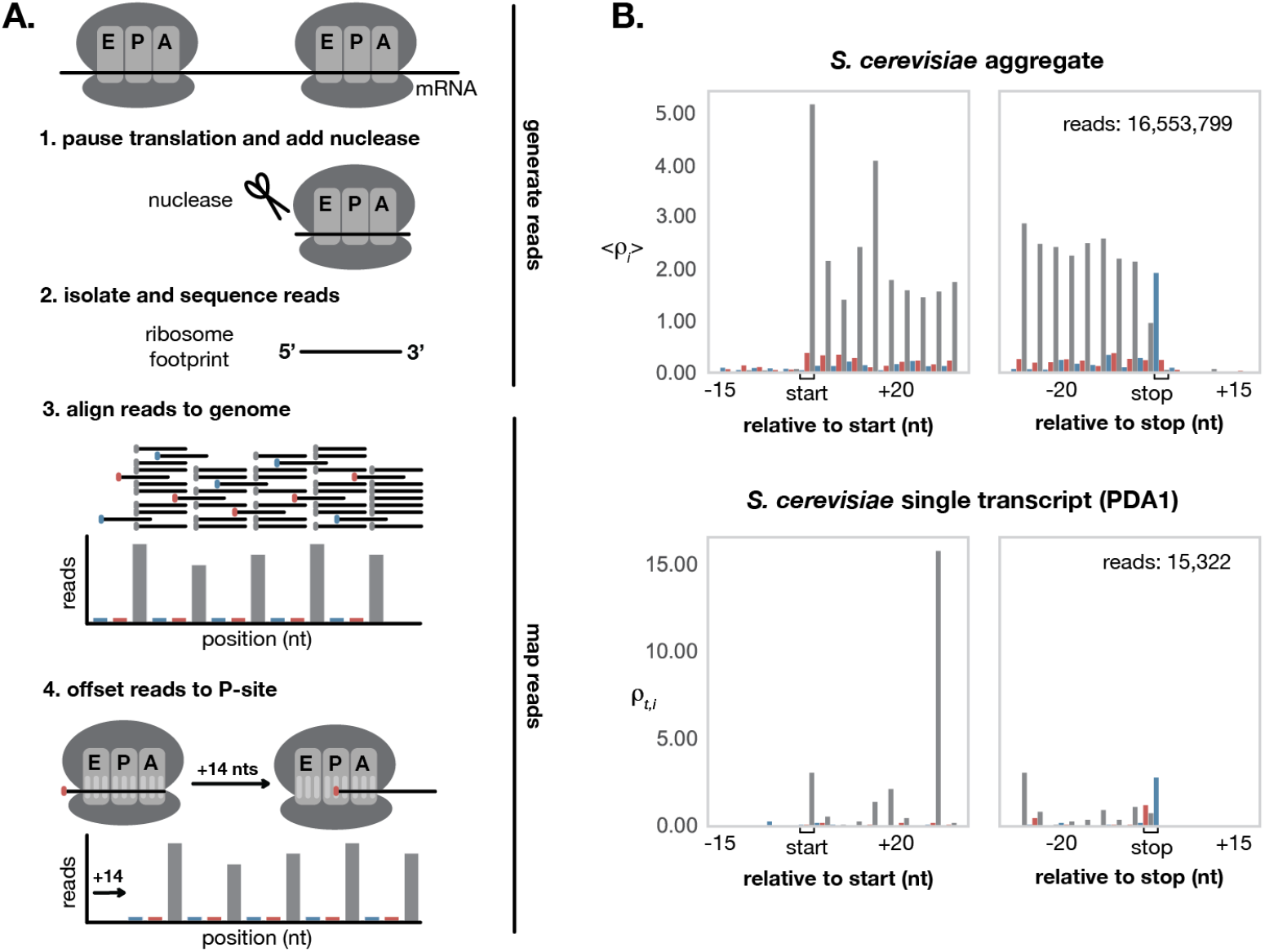
Processing ribo-seq data. (A) Ribo-seq involves experimentally generating reads (top), then computationally mapping sequenced reads to precise ribosome positions (bottom). Ideal data shows a clear triplet periodicity and can be offset to correspond to a position within the P-site of the ribosome by shifting the 5’ or 3’ end of all reads. (B) Realigned ribo-seq data from Wu et al [38] shows clear triplet periodicity in aggregate across all annotated ORFs in *S. cerevisiae* (top). This periodic pattern is not as obvious in individual ORFs (bottom). The ribo-seq signal shown here for PDA1, a housekeeping gene commonly used as a reference in gene expression studies, is representative of the signal observed across other genes in *S. cerevisiae*.

The first step performed by ORFeus is a data processing step to combine information from the input aligned ribosome profiling data, reference annotations, and reference genome sequence. During this data processing step, ORFeus associates each protein-coding transcript (5’UTR, ORF, and 3’UTR) to its aligned ribo-seq read counts and nucleotide sequence using the annotations and genome sequence. For bacteria, we split the genome into one “transcript” per protein-coding ORF (each “transcript” corresponds directly to one annotated ORF plus any annotated UTRs in the upstream/downstream intergenic regions); i.e. we ignore operon structure and analyze one ORF at a time. To control for variation in both length and expression across different transcripts, ORFeus converts input read counts to relative ribo-seq density, which we call *ρ*. The relative ribo-seq density 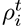 at position *i* of transcript *t* is calculated by normalizing the raw read counts at position *i* of transcript *t* by the mean read counts per position for transcript *t*: 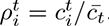. This per-transcript normalization ensures that ribo-seq density values are comparable across different transcripts, so the model can expect similar magnitude 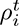 values within each translated ORF. Note, however that transcripts with especially long UTRs with no coverage will have lower 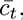 and thus higher 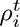 relative to transcripts with short or no UTRs. When aggregated across all transcripts, ideal ribo-seq density values should be near zero in the UTRs and exhibit clear periodicity within the ORFs (Fig. 1B).

### ORFeus algorithm

ORFeus is designed to detect both canonical and non-canonical ORFs from ribo-seq data (Fig. 2A). It uses a hidden Markov model (HMM) trained to recognize riboseq signal and nucleotide sequence features characteristic of translated ORFs. HMMs provide a probabilistic framework well-suited for handling noisy and sparse signals like ribo-seq data. [39] An HMM predicts the most probable path of hidden “states” that could generate the observed data. In our case, ORFeus takes as input information about the ribo-seq reads and nucleotide sequence at each position of a transcript and returns the most likely path of ORF or non-ORF states for that transcript.

**Fig. 2.**
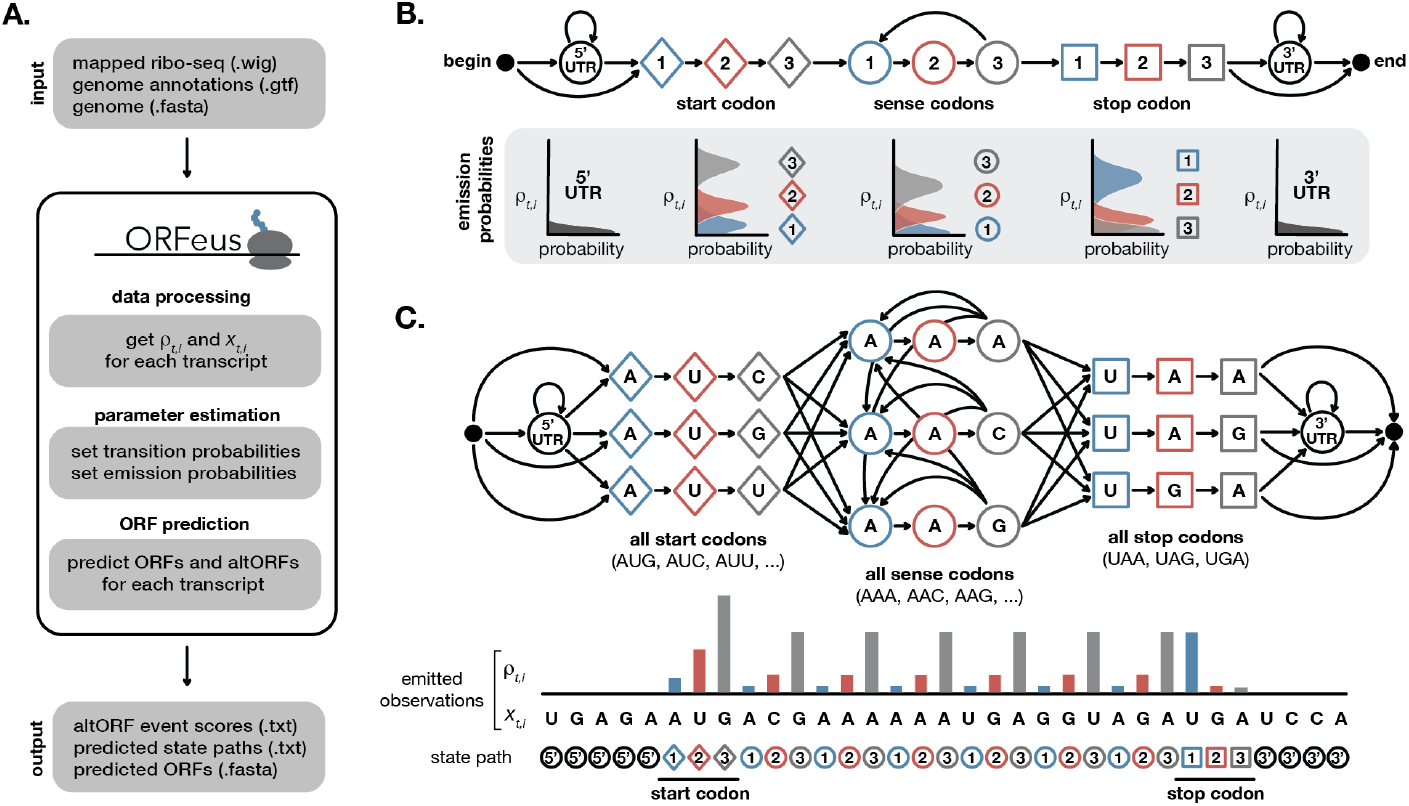
Inferring canonical ORFs from ribo-seq data. (A) ORFeus takes in mapped ribo-seq data, genome annotations, and genome sequences and returns predicted ORFs calculated using an HMM. (B) A simple HMM to model canonical translation includes states corresponding to the 5’UTR and 3’UTR as well as nucleotide 1 (blue), 2 (red), and 3 (grey) of each start codon (diamonds), sense codon (circles), and stop codon (squares). All nonzero transition probabilities are denoted with arrows. A schematic representation of the emission probabilities for each state are plotted below the corresponding states. (C) A more complex HMM includes states for each individual start, stop, and sense codon sequence. Idealized ribo-seq density for a transcript that undergoes canonical translation of a single ORF is depicted below the model. The input relative ribo-seq density 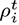 and nucleotide sequence 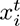 at each position inform the output state path prediction.

A simple HMM to model a canonical ORF includes eleven types of states: a 5’UTR state, a 3’UTR state, and states corresponding to nucleotide 1, 2, and 3 of each start codon, sense codon, and stop codon. The simple model (Fig. 2B) generates ribo-seq 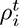 values for each position in the transcript. There is additional information in the nucleotide sequence of the transcript, including codon usage preferences and start/stop codon preferences. To accommodate this codon information in the HMM, we expand the model to also emit a nucleotide for each state. The resulting model includes separate trios of states for each possible start, sense, and stop codon. This model (Fig. 2C) generates two observed sequences: the ribo-seq values 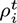 and the nucleotide sequence 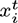. Valid start codons are determined by default from the annotations and genome sequence files, but can be altered by the user. This enables non-canonical start codon detection.

To model non-canonical ORFs, we add additional states and transitions (Fig. 3A). Our original goal was just to capture programmed ribosomal frameshifts. To model a +1 frameshift we added an *X*_1_ state to represent the nucleotide that is skipped over during translation. Translation can then resume in the +1 reading frame, continuing to a sense codon nucleotide 1 state (Fig. 3A blue arrows). We then chose to model a -1 frameshift as a +2 frameshift, since the resulting downstream frame should be equivalent and this is a reasonable approximation given the resolution of ribo-seq data. To shift the reading frame forward by two nucleotides (equivalent to back by one nucleotide), we added a second state *X*_2_ to represent the second nucleotide that is skipped over during a +2 frameshift (Fig. 3A red arrows).

**Fig. 3.**
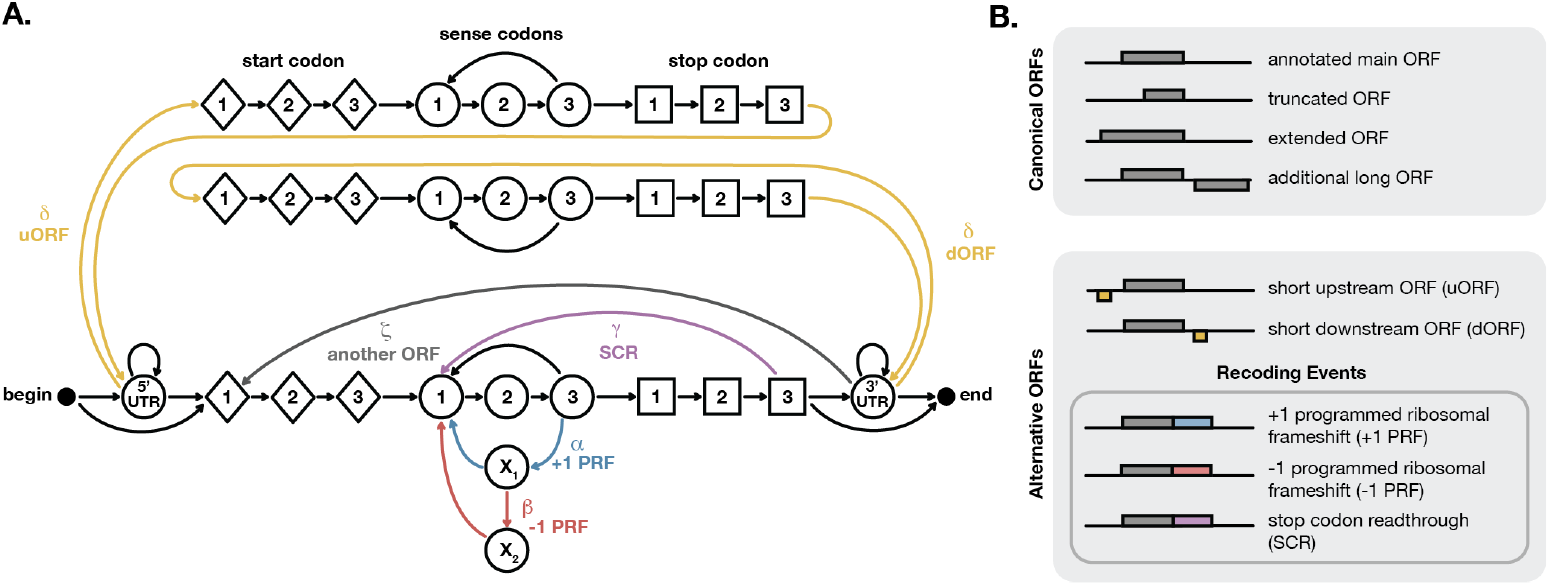
Inferring alternative ORFs from ribo-seq data. (A) A more complex HMM to model noncanonical translation includes additional states and transitions to represent upstream ORFs and downstream ORFs (yellow), programmed ribosomal frameshifting (blue and red), and stop codon read-through (purple). The probability of each type of event (denoted with a Greek letter) is set to reflect its expected frequency. (B) The full model can infer both canonical ORFs and alternative ORFs, including recoding events.

Since the HMM framework is very general, we can also add states and transitions to capture essentially all types of alternative translation events. We added stop codon readthrough by allowing movement from a stop codon directly back to a new sense codon (Fig. 3A purple arrow). To model short upstream and downstream ORFs, we added special start, sense, and stop codon states that can be accessed only from within the 5’UTR or 3’UTR respectively (Fig. 3A yellow arrows). These uORF and dORF states are included as separate states from the main ORF since their lengths are expected to be much shorter. Multiple long ORFs are included by allowing a transition from a 3’UTR into another main ORF (Fig. 3A grey arrow). The full model predicts several types of canonical and non-canonical translation (Fig. 3B).

#### Parameter estimation

After the ribo-seq data has been processed, ORFeus is trained on the input data set to recognize the signals for ORF and non-ORF states, with data-specific parameters. There are two sets of probabilities that must be estimated:

- **Transition probabilities** represent the probability of moving to state *k* from state *k −* 1. These are represented by arrows in Fig. 2B and 2C. For example, the probability of moving from a 5’UTR state into a start codon state is represented by the arrow from the 5’UTR state to the start nucleotide 1 state.
- **Emission probabilities** represent the probability of observing a specific ribo-seq value 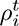 and the nucleotide sequence 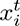 in state *k*. Different states have different expected ribo-seq density values. These are represented by rotated histograms below the state map in Fig. 2C. For example, we expect mostly 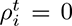 for positions in the 5’UTR state (dark grey histogram below the 5’UTR state), while we expect to observe higher 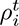 values more often for positions in start codon states (blue, red, and light grey histograms below the start codon). The nucleotide identity also factors into the emission probabilities for the expanded model shown in Fig. 2C, since only some nucleotides are allowable for certain states (e.g. UAA, UAG, and UGA for stop codons). Most states emit their designated nucleotide 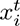 with probability 1.0.

#### Transition probabilities

Except for the few transitions discussed below, the transition probabilities are set to the observed frequency of each type of transition, based on the input transcriptome (generated from the annotations and genome sequence as described in data processing). For example, the probability of transitioning from the begin state into a 5’UTR is proportional to the frequency of annotated transcripts with 5’UTRs in the input transcriptome. The probability of remaining in a 5’UTR versus transitioning to a start codon is based on the mean length of annotated 5’UTRs. The probability of moving from the 5’UTR into state 1 of a particular start codon also depends on the relative frequency of this start codon across all annotated transcripts. Similarly, the probability of moving from state 3 of one sense codon to state 1 of the next sense codon is the relative usage of this next codon across all annotated transcripts. In this way, most of the model’s transitions are set to reflect features specific to the input transcriptome. All transition probabilities are enumerated in Table S1.

There are six transition probabilities that are set to the expected frequency of non-canonical translation events (denoted by greek letters in Fig. 3A). These six probabilities, which we call altORF event parameters, are set by hand. We have little *a priori* knowledge of the rate of multiple ORFs, stop codon readthrough, or programmed ribosomal frameshifting across transcriptomes, and we cannot estimate them from the annotations, so we set these values instead by optimizing correct identification of known events across our test data sets. We chose a single set of values that worked to identify known events across all of our test data sets. The probability of frameshifting per codon *α* = 10*^−^*^5^ is the total probability of a programmed ribosomal frameshift event, either in the +1 or -1 direction. The probability of -1 frameshift given that a frameshift occurs *β* = 0.5 represents the relative frequency of +1 and -1 frameshifting (thus we assume that both types of frameshifting are equally likely). The probability of stop codon readthrough per ORF *γ* = 10*^−^*^4^ is the total probability of translation past an in-frame stop codon, whether by sense codon incorporation or stop codon bypassing. The probability of short ORFs (including uORFs and dORFs) *δ* = 10*^−^*^3^ is the probability of observing a short ORF at each nucleotide in the 5’UTR or 3’UTR. The probability of multiple non-overlapping ORFs *ζ* = 10*^−^*^10^ is the probability of initiating another long ORF (including longer uORFs and dORFs) at each 3’UTR position downstream of the main ORF. Each of these parameters can be adjusted by the user to more accurately reflect the expected probabilities of alternative translation in the transcriptome of interest.

#### Emission probabilities

The emission probabilities are set to the probability of observing a particular nucleotide and a particular relative ribo-seq density 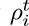 value in each state. The total emission probability is therefore the product of the nucleotide emission probability and the ribo-seq emission probability. The nucleotide emission probability is one over the number of possible nucleotides if the nucleotide is represented by this state, and zero otherwise. For example, the probability of observing A, T, C, or G in a UTR state is 1/4, since any valid nucleotide may be represented by these states (and we chose not to incorporate information about UTR nucleotide composition to avoid biasing our search against finding novel ORFs in annotated UTRs). In contrast, the probability of observing A in an ACG1 sense codon state is 1, since the first nucleotide state of an ACG codon must be an A. The ribo-seq emission probability is calculated for each state using the frequency of 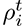 observed across all annotated protein-coding transcripts. For example, the probability of observing 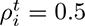 in an ACG1 state is set to the frequency of 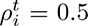 values across all first positions of annotated ACG sense codons. The ribo-seq emission histograms are binned to speed up downstream calculations, with 25 uniform bins spanning the range of observed 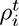 values in the input ribo-seq data.

Since each start, sense, and stop state represents a single codon, differences in signal that are sequence-specific can be picked up by the model. Nucleases used in ribosome profiling can exhibit nucleotide bias, preferentially cleaving after certain nucleotides leading to added noise. For example, RelE usually cleaves after the second nucleotide of the A-site, but prefers to cleave after a C nucleotide (and strongly avoids cleavage before a C nucleotide).[20] As a result, NNC codons are more often cleaved after the third nucleotide of the A-site instead, leading to a disruption in periodicity at these codons. [20] The codon-specific emissions in our model recognize this bias, expecting to see density at the third nucleotide in NNC codons and the second nucleotide in other codons. This strategy turns nuclease bias, which is a common source of noise in ribo-seq data, into signal that the model can use to inform reading frame prediction. This is useful in all cases except for especially small transcriptomes (such as viral or organellar transcriptomes). When there are too few transcripts to train emissions for each possible codon, we suggest pooling all codons during calculation of the emissions, which is an option available to the user.

The emissions for non-canonical ORF states are set after the canonical ORF emissions have been calculated. The emissions for uORF and dORF states are set to be the same as those for a canonical ORF. The emissions for the frameshift states *X*_1_ and *X*_2_ are naively set to the mean of the emissions for states 1, 2, and 3, since we have no prior knowledge about what these distributions should look like (since known frameshifts are rare).

After estimating all emission probabilities, we add a pseudocount of 10*^−^*^10^ to each ribo-seq emission probability bin and then re-normalize so that the total emission probability for each state is still one. This ensures that there is a non-zero probability of observing any possible ribo-seq value in each state. The advantage of this is that it allows the model to consider paths that go through unlikely (but not impossible) states for a given sequence and lets the model predict ORFs that contain ribo-seq values that were not observed in annotated ORFs for the given data set.

#### ORF prediction

After the parameters have been trained, ORFeus is ready to predict ORFs for individual transcripts. For each transcript, ORFeus returns the most probable state path. This indicates the positions and sequences of predicted canonical and non-canonical ORFs. We use the Viterbi algorithm [40] to compute the most probable state path for each transcript.

The output path tells us whether an altORF is inferred. The predicted state path for a +1 frameshift event includes state *X*_1_, which shifts the downstream frame forward by one nucleotide. The predicted state path for a -1 frameshift (or +2 frameshift) event includes states *X*_1_ and *X*_2_, which shifts the downstream frame forward by two nucleotides (which is is equivalent to shifting the downstream frame backward by one nucleotide). Similarly, the presence of uORF or dORF states indicates multiple ORFs are predicted and gives the exact inferred sequence of these additional ORFs. Finally, to indicate stop codon readthrough, the Viterbi path includes the transition from a stop state 3 back to sense state 1.

### Model Testing

#### Known altORFs

To test ORFeus, we ran the model on known examples of alternative translation. We used examples from multiple species to evaluate the method, which can be run on data from varied organisms and ribo-seq experimental protocols. We trained and ran the model on published data from *E. coli* (Hwang & Buskirk, 2017) [20], *S. cerevisiae* (Wu et al, 2019) [38], *D. melanogaster* embryos (Dunn et al, 2013) [16], *D. rerio* embryos (Bazzini et al, 2014) [31], and SARS-CoV-2 infected *C. sabaeus* Vero E6 cells (Finkel et al, 2021) [41]. Known examples of altORFs in these species were used to tune the altORF event parameters to a single *α*, *β*, *γ*, *δ*, and *ζ* value that can be used across all tested data sets.

With these altORF event parameters, ORFeus correctly identifies well-characterized examples of alternative translation, including: a +1 frameshift in *E. coli* prfB (Fig. 4A), a -1 frameshift in SARS-CoV-2 ORF1ab (Fig. 4B), stop codon readthrough in *D. melanogaster* headcase (hdc) (Fig. 4C), uORFs upstream of *S. serevisiae* GCN4 (Fig. 4D), and a dORF downstream of *D. rerio* rrm1 (Fig. 4E). *D. melanogaster* hdc has an abundance of ribo-seq signal in the 5’UTR. With our chosen default parameters, ORFeus does not predict any uORFs to account for this signal, but with slightly different parameter choices it does. We were unable to distinguish whether this signal, which was also recognized by Dunn et al [16]), represents a real translation signal or some sort of artifact.

**Fig. 4.**
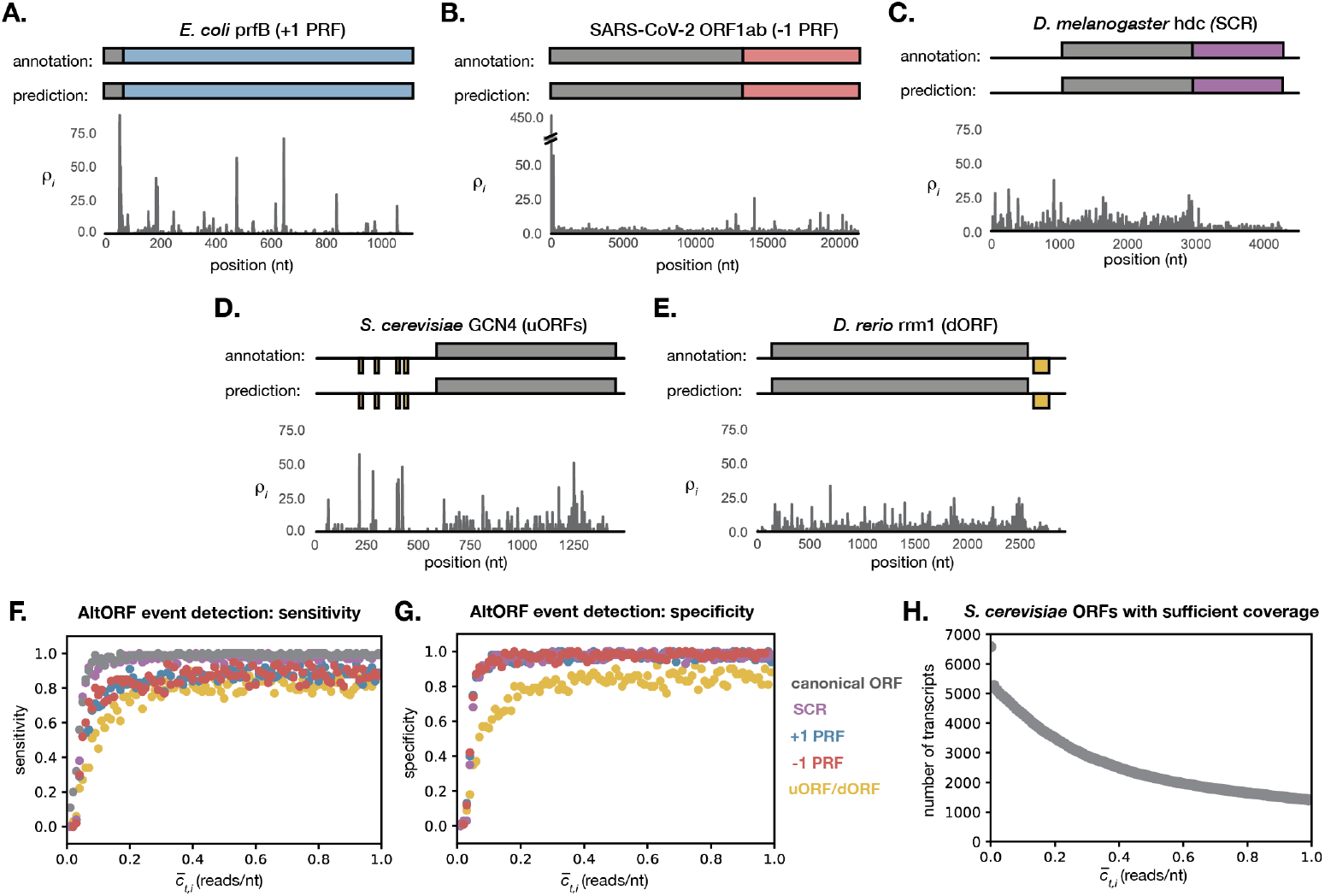
Model performance on known and simulated altORFs. (A-E) Well-characterized examples of alternative translation events are correctly inferred by the model. The annotated and predicted ORF are shown schematically above the real 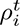 signal for each transcript. Sensitivity (F) and specificity (G) are shown for sequences simulated using parameters estimated from the Wu et al *S. cerevisiae* data set. [38] Each point represents the mean sensitivity or specificity over 100 simulated sequences with the given mean ribo-seq coverage. (H) The number of ORFs with at least the given ribo-seq coverage drops off rapidly as the coverage threshold increases. The data shown are for the Wu et al *S. cerevisiae* data set. [38]

These examples show that ORFeus is capable of detecting real altORF events. A subset of these events can be detected in data sets lacking exceptionally clear triplet periodicity (stop codon readthrough in *D. melanogaster*). However, the number of known cases is anecdotal, and it is important to note that the altORF event parameters used to detect these events were optimized for good prediction on these test cases themselves.

#### Simulated altORFs

To evaluate the performance of ORFeus across a transcriptome, we want to estimate sensitivity and specificity of altORF event detection. Since there are a limited number of known examples of non-canonical translation, we cannot systematically evaluate the model’s performance on real data. Instead, we devised a method to simulate altORFs from the model. Since an HMM is a generative model, we have the ability to simulate altORFs using the transitions and emissions. We create a state path using the transition probabilities, then select the nucleotide sequence and relative ribo-seq density 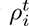 values using the emission probabilities for each state.

Sampling from an HMM to generate a simulated ORF is trivial. Here though, we specifically want to simulate altORFs, which include a transition into a specific set of altORF states. Since the probability of transitioning into any given altORF states is rare, we would have to simulate thousands of ORFs from the HMM to generate a single altORF by chance. So instead, we sample *conditional* on the state path including a transition into an altORF event. We use a sampling method that works outward from the desired altORF event. The forward sampling direction follows the standard HMM sequence generation method, but the backward direction depends on a transition matrix inversion (Implementation).

Sequences can also be simulated to have a target mean ribo-seq coverage level 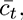 To do this, we first estimate the total number of reads *N_t_* that should be assigned to the transcript to generate this mean coverage level: 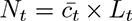 where *L_t_* is the total length of the transcript in nucleotides. Reads are then assigned to each position *i* in the transcript with probability proportional to 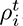.

The sensitivity and specificity for altORF detection at different ribo-seq coverage levels were estimated using sequences simulated by this approach (Fig. 4G and 4H). The model was run on the simulated sequence of nucleotides and relative ribo-seq density values for ORFs of various coverage levels. If an altORF event was simulated and then correctly predicted within *±*10 nucleotides of the true simulated location, the result was considered a true positive. If no altORF event was simulated, but an altORF event was predicted, the event was considered a false positive.

The ribo-seq coverage level needed for accurate altORF detection will vary by data set. Before running ORFeus on a given ribo-seq data set, we recommend running the sensitivity and specificity simulations (included in the ORFeus code) to determine a coverage threshold that yields appropriate performance (Fig. 4F and 4G). ORFeus should only be run on transcripts with this ribo-seq coverage or higher. The number of transcripts in a given data set with the necessary coverage level will depend on the coverage threshold chosen (Fig. 4H).

## Discussion

### Limitations

ORFeus provides no mechanistic insights about *how* a predicted non-canonical translation event occurs, and is therefore limited in the scope of what types of events can be called. For example, we are unable to distinguish between alternate forms of stop codon readthrough. ORFeus can only identify that there is translation immediately past a stop codon, not whether it is due to selenocysteine incorporation, near-cognate tRNA decoding, or bypassing of the stop codon altogether. All of these events result in downstream in-frame translation past a potential stop codon and are indistinguishable from each other at the current resolution of ribosome profiling data. Similarly, we cannot distinguish a +1 programmed ribosomal frameshift from a -2 frameshift (or a -1 frameshift from a +2 frameshift), which could result in the same downstream riboseq signal. In fact, we even model a -1 frameshift event as a +2 frameshift event given the lack of resolution to distinguish between their output, because it is convenient in the structure of the HMM.

Importantly, we limit ourselves to identifying events that could result in pronounced signal changes. For example, we search only for uORFs and dORFs that are translated at a rate comparable to the main ORF, because setting a lower expected signal is more likely to pick up noise. ORFeus is also only capable of predicting frameshifts that generate C-terminally extended protein products (e.g. *E. coli prfB* +1 frameshift), since it requires that there be ribo-seq signal in a new frame downstream of the frameshift site. It is unable to identify frameshifts that result in early termination, where little or no sequence is translated in the alternate frame after the frameshift site (e.g. *E. coli dnaX* -1 frameshift).

### Applications

Running ORFeus on published or new whole-transcriptome ribo-seq data sets may uncover previously overlooked or unseen translation products. Even in well-annotated genomes, it is likely that alternative protein sequences are translated and have escaped detection by proteomics due to short length or low abundance. For organisms with less well-annotated genomes, ORFeus provides an opportunity to search for both canonical ORFs and non-canonical translation products.

## Conclusions

We developed an HMM-based model for inferring both canonically and non-canonically translated ORFs from ribo-seq data. With the ability to detect non-standard translation, ORFeus is a powerful, general-purpose tool for uncovering potential novel protein products and expanding our knowledge of translation across different organisms.

## Methods

### Transcript annotations

Genome sequence and annotations were downloaded for *E. coli*, *S. cerevisiae D. melanogaster*, *D. rerio*, and SARS-CoV-2 from Ensembl [42] (Table 1). Since alternative translation can lead to translation outside of annotated ORFs, it was important to have complete UTR annotations for all species. We updated the annotations and transcript sequences for *E. coli* to include UTRs from RegulonDB [43] and *S. cerevisiae* to include UTRs from Nagalakshmi et al [44].

**Table 1.** Data sources used in this study. P-site position indicates the nucleotide of the P-site where most reads map, after the P-site offset is applied. Aindicates that the periodicity in the data is not clear enough to determine the exact nucleotide within the P-site.

| Organism | Genome Version | SRA Accession(s) | Read Lengths | P-site Offset | P-site Position | Ribo-Seq Data Reference |
| --- | --- | --- | --- | --- | --- | --- |
| Coronavirus (SARS-CoV-2) | ASM985889v3 | SRR12216748-50 | 28-30nt | 5'+14nt | 3rd nt | Finkel et al, 2021 |
| *E. coli* (K12, MG1655) | ASM584v2 | SRR4023281 | 20-40nt | 3'-3nt | 3rd nt | Hwang & Buskirk, 2017 |
| *S. cerevisiae* (S288C) | R64-1-1 | SRR7241903-04 | 28nt | 5'+14nt | 3rd nt | Wu et al, 2019 |
| *D. melanogaster* (embryos) | BDGP6.28 | SRR942868-71,74-79 | 20-40nt | 5'+16nt | ? | Dunn et al, 2013 |
| *D. rerio* (embryos) | GRCz11 | SRR1062294-302 | 28-29nt | 5'+12nt | 1st nt | Bazzini & Johnstone et al, 2014 |

### Ribo-seq data

In order to infer ORFs, ORFeus requires information about the relative density of elongating ribosomes at each position of annotated transcripts. We downloaded raw ribo-seq read data for *E. coli* [20], *S. cerevisiae* [38], *D. melanogaster* embryos [16], *D. rerio* embryos [31], and SARS-CoV-2 infected *C. sabaeus* Vero E6 cells [41] (Table 1).

We realigned all data sets, since aligned data was not available for many of the studies. This also allowed us to align to the most recent genome version and to offset reads to align to the P-site, which is necessary for correct analysis with ORFeus. Wherever possible, we attempted to replicate the methods used to align the data in the original reference. Each of the raw ribo-seq libraries was processed by: (i) trimming adapters and low quality bases (phred score below 20) with Cutadapt v1.8.1 (Martin, 2011); (ii) removing reads mapping to ladder sequences and organism-specific non-coding RNAs from Ensembl (Howe et al, 2019); (iii) aligning remaining reads uniquely to reference genome sequences from Ensembl; (iv) filtering reads by length and offsetting the reads so they correspond to the P-site of the ribosome using Shoelaces [21].

Reads from SARS-CoV-2, *E. coli*, and *S. cerevisiae* were aligned uniquely using Bowtie1 v1.1.1 (-v 2 -y -m 1 -a –best –strata) [45], since few or no introns are present. Reads from *D. melanogaster* and *D. rerio* were aligned using the splice-aware aligner STAR v2.7.0 (–outSAMmultNmax 1 –outFilterMultimapNmax -1 –outFilterMismatchNmax 2) [46] and uniquely mapped reads were identified with Samtools v1.10 (view -h -q 255) [47]. For the offset to the P-site, the exact nucleotide position within the P-site was chosen separately for each data set. We selected the P-site position that resulted in any distinct start or stop codon signals being mapped to within the start or stop codon respectively, since distinct start and stop signals are modeled by ORFeus. For example, for the *S. cerevisiae* data [38], we used an offset of 14 nucleotides from the 5’ end (which corresponds to the 3rd nucleotide of the P-site) to ensure the distinct stop codon peak mapped within the stop codon (Fig. 1B). Read lengths and offsets selected for each experiment are shown in Table 1.

### Parameter estimation

The altORF event parameters were set to the following values: *α* = 10*^−^*^5^, *β* = 0.5, *γ* = 10*^−^*^4^, *δ* = 10*^−^*^3^, *ζ* = 10*^−^*^10^. All other parameters in the model were estimated separately for each ribo-seq data set and corresponding annotations file. Canonical start (AUG), stop (UAA, UAG, UGA), and sense codons were used. Transition probabilities between codons were set to the frequency of each codon in the annotated transcripts. Mean uORF and dORF lengths were set to 50 nts. Mean main ORF lengths were set to the mean ORF length of all annotated transcripts for all data sets.

### Sequence simulation

Sequences were simulated by generating a valid state path from the model (using the transition probabilities), then generating valid ribo-seq and nucleotide emissions from each state (using the emission probabilities). For calculation of sensitivity and specificity for rare non-canonical events, sequences were simulated starting at the rare event state(s) and extended in either direction: continuing forward until the end state and backward until the begin state. This was done to ensure the event would be present in the simulated sequence, despite it rarely occurring in normal simulation.

The transition probabilities were used to calculate paths forward during simulation, and the reverse transition probabilities were used to calculate paths backward. Reverse transitions are calculated according to equation (1), as outlined by Solow & Smith[48].

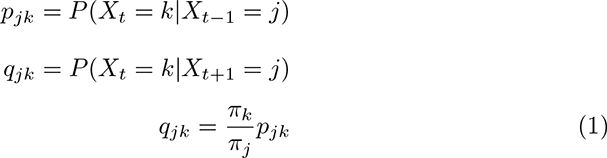

This computation requires that the model meet the conditions of reversibility: stationary (transition matrix doesn’t change over time), irreducible (each state can eventually be reached from every other state), positive recurrent (expected return time to each state is finite), and aperiodic (stating in each state, there is no regular period at which the state cannot be reached). These conditions are met for the case where all states are accessible (i.e. *α*, *β*, *γ*, *δ*, and *ζ* are all nonzero). *π* is then the stationary distribution, which is computed by finding the eigenvector for the transpose of the transition probability matrix corresponding to the eigenvalue *λ* = 1.

### Coverage threshold

The minimum ribo-seq coverage needed to accurately infer altORFs was estimated using simulated sequences. We used the model to generate one hundred sequences per each mean coverage value from 0.01 footprints per nucleotide to 1.0 footprints per nucleotide, in steps of 0.01. To estimate sensitivity, we ran ORFeus on each simulated sequence and determined whether the output Viterbi path contained the correct noncanonical translation event (starting and ending within *±*10 nucleotides of the true simulated positions). For example, a programmed ribosomal frameshift was considered correctly inferred if it was up to 10 nucleotides upstream or downstream of the true simulated frameshift site. Similarly, a uORF, dORF, or canonical ORF was considered considered correctly inferred if its start and stop codons were within *±*10 nucleotides of the simulated start and stop codons respectively. To estimate specificity, we ran ORFeus on sequences simulated without any altORF events and determined whether the output Viterbi path contained any non-canonical translation event (at any position in the sequence).

## Abbreviations

ORF: open reading frame
altORF: alternative ORF
uORF: upstream ORF
dORF: downstream ORF
SCR: stop codon readthrough
PRF: programmed ribosomal frameshift

## Declarations

### Ethics approval and consent to participate

Not applicable

### Consent for publication

Not applicable

### Availability of data and materials

All code and data necessary to reproduce and extend the work is freely available at http://eddylab.org/publications/#RichardsonEddy23 and on GitHub at https://github.com/morichardson/ORFeus/releases/tag/v1.0.0.

### Authors’ contributions

MOR conceived, designed, and carried out the work under the mentorship of SRE. MOR drafted the manuscript and both authors contributed to and approved the final version.

## Acknowledgments

We thank Elena Rivas, Andrew Murray, and members of the Eddy lab for discussions and for feedback on the manuscript. Computations in this paper were run on the FASRC Cannon cluster supported by the FAS Division of Science Research Computing Group at Harvard University.

