## Supplemental Table 1 for "ORFeus: A Computational Method to Detect Programmed Ribosomal Frameshifts and Other Non-Canonical Translation Events"

| | **5'UTR** | **start$_1$** | **start$_2$** | **start$_3$** | **sense$_1$** | **sense$_2$** | **sense$_3$** | **stop$_1$** | **stop$_2$** | **stop$_3$** | **3'UTR** | **X$_1$** | **X$_2$** | **uORF/ dORF start$_1$** | **uORF/ dORF start$_2$** | **uORF/ dORF start$_3$** | **uORF/ dORF sense$_1$** | **uORF/ dORF sense$_2$** | **uORF/ dORF sense$_3$** | **uORF/ dORF stop$_1$** | **uORF/ dORF stop$_2$** | **uORF/ dORF stop$_3$** | **end** |
|---|---|---|---|---|---|---|---|---|---|---|---|---|---|---|---|---|---|---|---|---|---|---|---|
| **begin** | $f_{5'}$ | $1-f_{5'}$ | | | | | | | | | | | | | | | | | | | | | |
| **5'UTR** | $\frac{L_{5'}-1}{L_{5'}}(1-\delta)$ | $(1-f_{5'})f_c$ | | | | | | | | | | | | $\delta f_c$ (uORF only) | | | | | | | | | |
| **start$_1$** | | | 1 | | | | | | | | | | | | | | | | | | | | |
| **start$_2$** | | | | 1 | | | | | | | | | | | | | | | | | | | |
| **start$_3$** | | | | | $f_c$ | | | | | | | | | | | | | | | | | | |
| **sense$_1$** | | | | | | 1 | | | | | | | | | | | | | | | | | |
| **sense$_2$** | | | | | | | 1 | | | | | | | | | | | | | | | | |
| **sense$_3$** | | | | | $\frac{L_{ORF}-1}{L_{ORF}}f_c(1-\alpha)$ | | | $\frac{1}{L_{ORF}}f_c(1-\alpha)$ | | | | $\alpha$ | | | | | | | | | | | |
| **stop$_1$** | | | | | | | | | 1 | | | | | | | | | | | | | | |
| **stop$_2$** | | | | | | | | | | 1 | | | | | | | | | | | | | |
| **stop$_3$** | | | | | $\gamma f_c$ | | | | | | $f_{3'}(1-\gamma)$ | | | | | | | | | | | | $(1-f_{3'})(1-\gamma)$ |
| **3'UTR** | | $\zeta f_c$ | | | | | | | | | $\frac{L_{3'}-1}{L_{3'}}f_{3'}(1-\delta)(1-\zeta)$ | | | $\delta f_c$ (dORF only) | | | | | | | | | $\frac{1}{L_{3'}}(1-\delta)(1-\zeta)$ |
| **X$_1$** | | | | | $f_c(1-\beta)$ | | | | | | | | $\beta$ | | | | | | | | | | |
| **X$_2$** | | | | | $f_c$ | | | | | | | | | | | | | | | | | | |
| **uORF/dORF start$_1$** | | | | | | | | | | | | | | | 1 | | | | | | | | |
| **uORF/dORF start$_2$** | | | | | | | | | | | | | | | | 1 | | | | | | | |
| **uORF/dORF start$_3$** | | | | | | | | | | | | | | | | | $f_c$ | | | | | | |
| **uORF/dORF sense$_1$** | | | | | | | | | | | | | | | | | | 1 | | | | | |
| **uORF/dORF sense$_2$** | | | | | | | | | | | | | | | | | | | 1 | | | | |
| **uORF/dORF sense$_3$** | | | | | | | | | | | | | | | | | $\frac{L_{sORF}-1}{L_{sORF}}f_c$ | | | $\frac{1}{L_{sORF}}f_c$ | | | |
| **uORF/dORF stop$_1$** | | | | | | | | | | | | | | | | | | | | | 1 | | |
| **uORF/dORF stop$_2$** | | | | | | | | | | | | | | | | | | | | | | 1 | |
| **uORF/dORF stop$_3$** | 1 (uORF only) | | | | | | | | | | 1 (dORF only) | | | | | | | | | | | | |

**Table S1:** Complete transition probability matrix for ORFeus. The row label represents the current state, and the column label represents the next state in the path. For example, the probability of transitioning from the begin state to the 5'UTR is shown in the first cell. All blank cells have a transition probability of zero, indicating that there is no probability of transitioning between the corresponding states in the table. The cells containing values each correspond to a transition arrow in Fig. 3A. $L_{5'}$ is the expected length of a 5'UTR in nucleotides; $L_{3'}$ is the expected length of a 3'UTR in nucleotides; $L_{ORF}$ is the expected length of a canonical ORF; $L_{sORF}$ is the expected length of a short uORF or dORF. $f_c$ is the frequency of the next codon in the state path. For example, $f_c$ at the transition from 5'UTR to start$_1$ is the frequency of the next start codon across all start codons (e.g. fraction of start codons that are AUG). Similarly, $f_c$ at the transition from 5'UTR to start$_3$ to sense$_1$ is the frequency of the next sense codon across all sense codons. Note that uORFs and dORFs have the same set of transition probabilities, except that uORFs can begin and end only in a 5'UTR (indicated by "uORF only"), while dORFs can begin and end only in a 3'UTR (indicated by "dORF only").
